# Docking Molecular analysis of potential Drug Paritaprevir against Mycobacterium tuberculosis (Mtb)

**DOI:** 10.1101/2021.07.09.451739

**Authors:** Ivan Vito Ferrari, Paolo Patrizio

## Abstract

**Background:** Mycobacterium tuberculosis (Mtb) is the causative agent of tuberculosis, which kills 1.8 million annually. This is an infectious disease generally affects the lungs, but can also affect other parts of the body. Mtb RNA polymerase (RNAP) is the target of the first-line antituberculosis drug Rifampin (Rif). We report first time a Potential Drug Paritaprevir against with severe infectious disease, by in Silico approach, using Autodock Vina and Autodock 4 (or MGL Tool), estimated with Pyrx and AMDock Software, calculating three different important parameters: Binding Affinity (kcal/mol), estimated Ki (in nM units) and Ligand Efficiency (L.E. in kcal/mol). After a selective analysis of over 1000 drugs, processed with Pyrx (a Virtual Screening software for Computational Drug Discovery) in the Ligand Binding site pocket of the protein (ID PDB 5UHB chain C,DNA-directed RNA polymerase subunit beta), we noticed high values of these 3 parameters mentioned above of Paritaprevir, concluding that it could be an excellent candidate drug for this type of infection. Indeed, from the results of Autodock Vina and Autodock 4 (or Autodock 4.2), implemented with lamarckian genetic algorithm, LGA, trough AMDock Software, This oral drug, approved by FDA in 2014, both by Autodock Vina and Autodock Vina 4 has excellent Binding affinity value, ca. −10.00 kcal/mol, a Ki value 40 nM and Ligand efficiency ca −0.15 kcal/mol. These results are comparable to the drug crystallized in the above-mentioned protein, currently used against TBC.

## 1. Introduction

Mycobacterium tuberculosis (Mtb) is the causative agent of tuberculosis, which kills 1.8 million annually. Mtb RNA polymerase (RNAP) is the target of the first-line antituberculosis drug Rifampin (Rif) [1]. Most infections show no symptoms, in which case it is known as latent tuberculosis. The classic symptoms of active TB are a chronic cough with blood-containing mucus, fever, night sweats, and weight loss [2]. A total of 1.4 million people died from TB in 2019 (including 208 000 people with HIV). Worldwide, TBC is one of the top 10 causes of death and the leading cause from a single infectious agent (above HIV/AIDS) [2]. In this short communication, we investigated about 1000 drugs, through In Silico Docking approach, downloaded from Pubchem Database (https://pubchem.ncbi.nlm.nih.gov/). Molecular docking, has become a powerful tool for drug discovery and optimization. General speaking, there are several protein-ligand docking programs currently available. This bioinformatics methodology, is one of the most frequently used methods in structure-based drug design, due to its ability to predict the binding-conformation of small molecule ligands to the appropriate target (or receptor) binding site. In this paper particular attention, we have focused on Autodock Vina, estimated with Pyrx software, a simple Virtual Screening library software for Computational Drug Discovery (https://pyrx.sourceforge.io/), based on prediction the binding orientation and affinity of a ligand [ 3] and that of Autodock 4, opened by AMDOCK software, (a versatile graphical tool for assisting molecular docking with Autodock Vina and Autodock4, characterized to be automated docking of ligand to macromolecule by Lamarckian Genetic Algorithm and Empirical Free Energy Scoring Function [ 4]. Both Scoring Functions, were supplied by The Scripps Research Institute [ 3–4].

## 1. Materials and methods

### 2.1 Protein and Ligand Preparation before docking

5UHB [Crystal structure of Mycobacterium tuberculosis transcription initiation complex in complex with Rifampin] was prepared manually using different software, before molecular docking analysis. The first step, was downloaded from Protein Data Bank, https://www.rcsb.org/structure/5UHB) and save in pdb format. The second step were removed all unnecessary docking chains. In fact, only the C chain (DNA-directed RNA polymerase subunit beta) has been maintained and re-saved in pdb format. (See below figure 1) Next, were the removal of ligands and water molecules crystallized using Chimera software [ 5]. Later, polar hydrogens and Kollmann charges were added with Mgl Tool, (or called AutoDockTools, a software developed at the Molecular Graphics Laboratory (MGL) [6] As a last step they were added to the protein, any missing amino acids and the whole protein was minimized with the Swiss PDB Viewer Software [ 7]. In this way the protein was saved in pdb format and ready for docking, through Autodock Vina, estimated with a soft virtual screening library called Pyrx.On the other hand for Ligand Preparation, the first step, was to separate the crystallized drug from its protein, manually add all the hydrogens and their charges (with the MGL Tool software) and minimize it with MMFF94 force field [8], opened Pyrx software (https://pyrx.sourceforge.io/). This crystallized drug, Rifampin (RFP) was docked in the same binding area in order to accurately evaluate its Binding Affinity Vina Score with its protein and if there is overlapping of the drug Rifampin, between its crystallized version and in its docking version. (See below figure 2) In the area of the drug crystallized with the protein’s C chain, more than 1000 dockings were carried out, through Autodock Vina, estimated with Pyrx software, estimating their Binding Affinity (kcal/mol).

**Fig 1.**
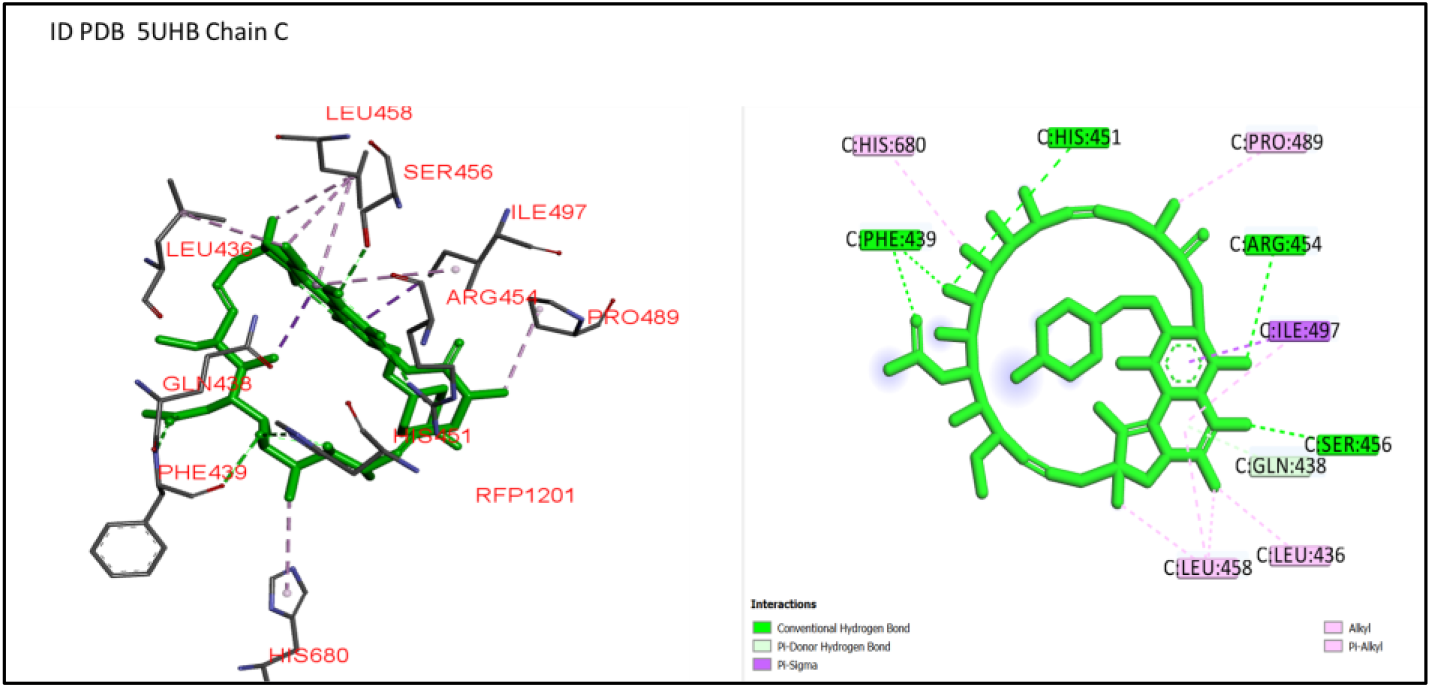
ID PDB 5UHB Chain C (DNA-directed RNA polymerase subunit beta), Comparison 3D Structure Cristalized and 2D Plot Diagram Cristalized Rifampin, In Ligand Binding Site pocket by Discovery Studio Biovia Software [9]

**Fig 2.**
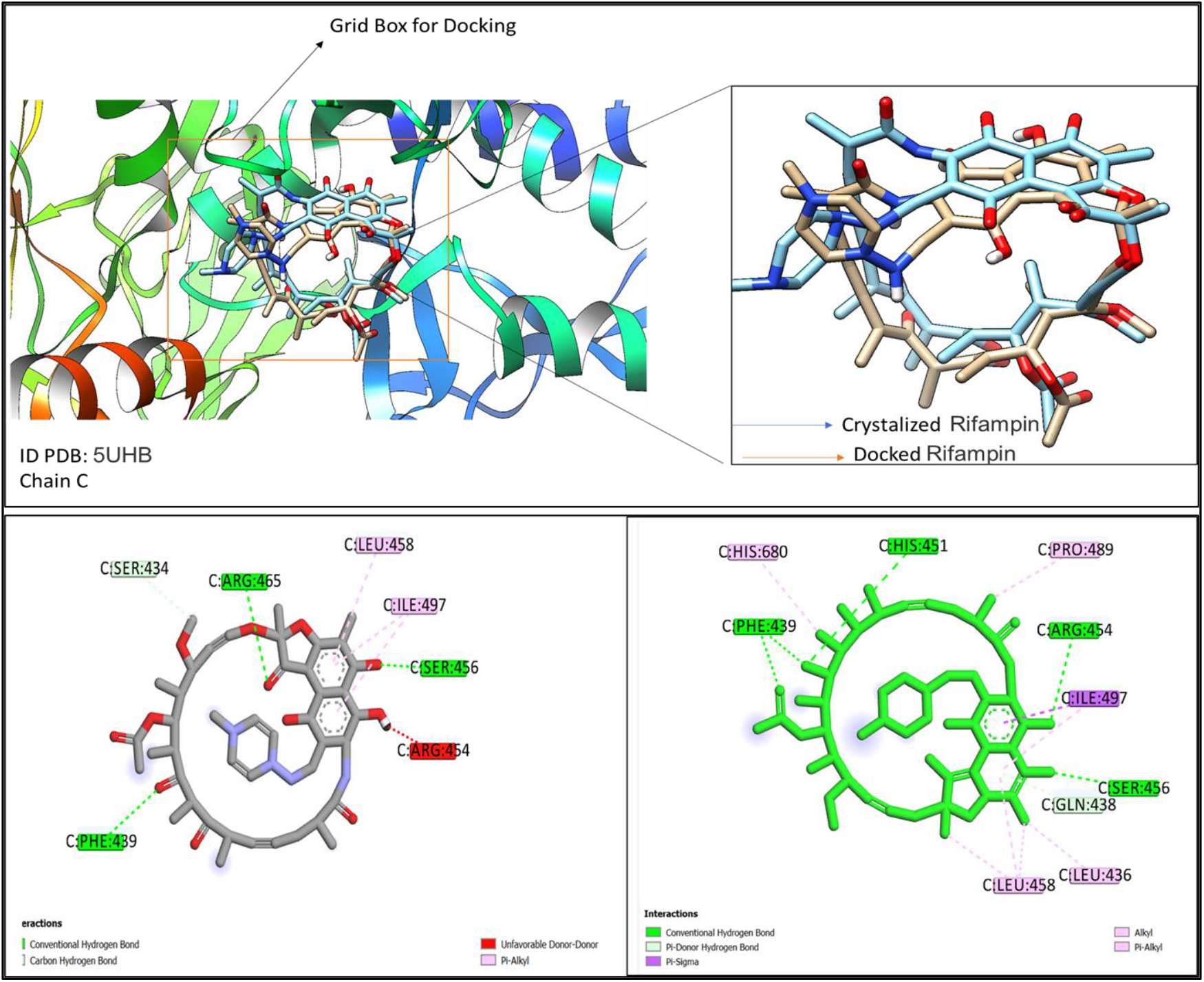
Comparison 3D Structure and 2D-Diagram, In Ligand Binding Site pocket preparated by Discovery Studio Biovia, [9] and Chimera Software [5] in ID PDB 5UHB Chain C protein (DNA-directed RNA polymerase subunit beta) complexed with crystalized Rifampin with docked Rifampin.

Parameters Grid Box for Docking in Ligand Binding Site Pocket for Repeatability Binding Affinity by AMDock Software calculated with Autodock Vina and Autodock Vina:

- ***ID PDB 5UHB Chain:** Center X (= 163.84); Centre Y (=162); Centre Z (=20); Dimensions (Angstrom) (Å) X, Y, Z [= 17=,17, =17]; exhaustiveness = 8*

## 3 Discussion and Results

We report first time a Potential Drug Paritaprevir, against with severe infectious disease TBC, by in Silico approach, using Autodock Vina and Autodock 4 (or MGL Tool), estimated with Pyrx and AMDock Software, calculating three different important parameters: Binding Affinity (kcal/mol), estimated Ki (in nM units) and Ligand Efficiency (L.E. in kcal/mol). After a selective analysis of over 1000 drugs, processed with Pyrx (a Virtual Screening software for Computational Drug Discovery) in the Ligand Binding site pocket of the protein (ID PDB 5UHB chain C, DNA-directed RNA polymerase subunit beta), we noticed high values of these 3 parameters mentioned above of Paritaprevir, concluding that it could be an excellent candidate drug for this type of infection. (See below figure 3) Indeed, from the results of Autodock Vina and Autodock Vina 4 (or Autodock 4.2), implemented with Lamarckian genetic algorithm, LGA, trough AMDock Software, This oral drug, approved by FDA in 2014, both by Autodock Vina and Autodock 4 has excellent Binding affinity value, ca. −10.00 kcal/mol, a Ki value 40 nM and Ligand efficiency ca −0.15 kcal/mol. These results are comparable to the drug crystallized in the above-mentioned protein, currently used against TBC. (See below figure 3)

**Fig.3.**
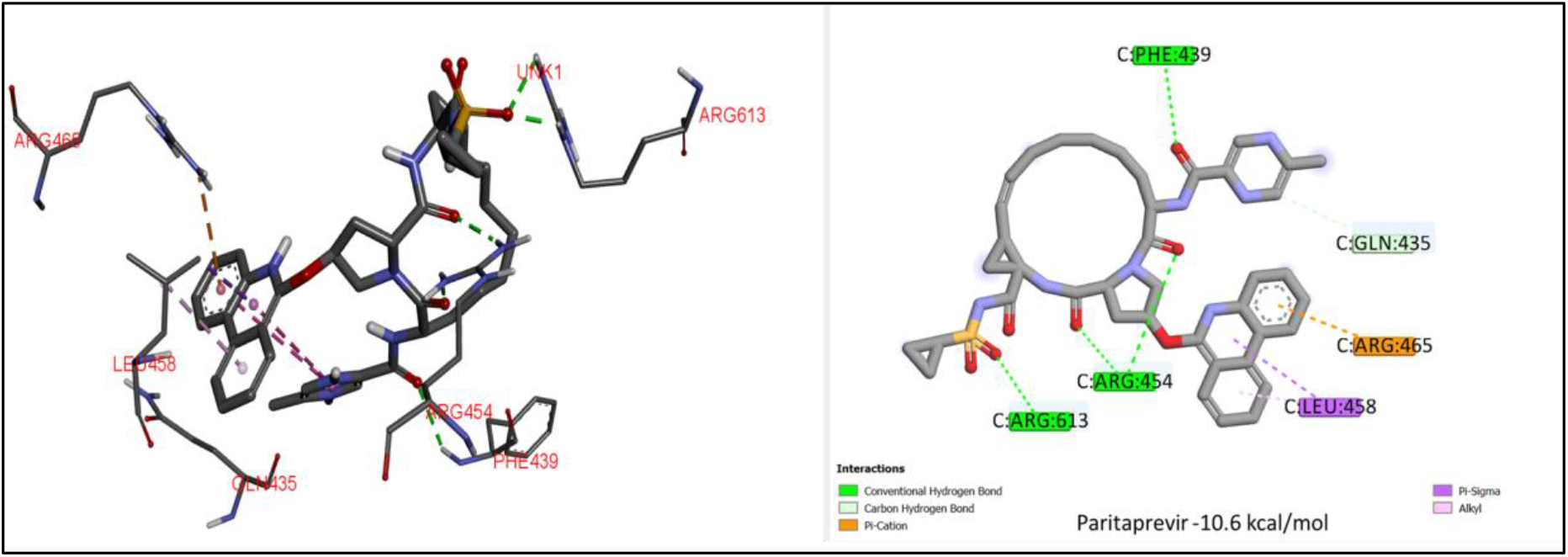
ID PDB 5UHB Chain C protein (DNA-directed RNA polymerase subunit beta: comparison 3D Structure Docked and 2D Plot Diagram Docked Paritaprevir In Ligand Binding Site pocket by Discovery Studio Biovia Visualizer Software [9]

**Tab.1.** Comparison results of Cristalized Drug Rifampin and proposed drug Paritaprevir calculated AutoDock Vina and AutoDock 4 (or MGL Tool), estimated by AMDock Software [4]

| DRUG | Scoring function | Binding affinity (kcal/mol) | Estimated Ki nM | Ligand efficiency (kcal/mol) |
| --- | --- | --- | --- | --- |
| Cristalized Rifampin | Autodock Vina | -8,9 | 299,41 | -0,15 |
| Paritaprevir | Autodock Vina | -10 | 46,77 | -0,18 |
| Cristalized Rifampin | Autodock 4 (MGL TOOL) | -9,87 | 58,24 | -0,17 |
| Paritaprevir | Autodock 4 (MGL TOOL) | -10,08 | 40,86 | -0,18 |

## 4. Conclusion

We report first time a Potential Drug Paritaprevir against with severe infectious disease, by in Silico approach, using Autodock Vina and Autodock 4 (or MGL Tool), estimated with Pyrx and AMDock Software, calculating three different important parameters: Binding Affinity (kcal/mol), estimated Ki (in nM units) and Ligand Efficiency (L.E. in kcal/mol). After a selective analysis of over 1000 drugs, processed with Pyrx (a Virtual Screening software for Computational Drug Discovery) in the Ligand Binding site pocket of the protein (ID PDB 5UHB chain C,DNA-directed RNA polymerase subunit beta), we noticed high values of these 3 parameters mentioned above of Paritaprevir, concluding that it could be an excellent candidate drug for this type of infection. Indeed, from the results of Autodock Vina and Autodock 4 (or Autodock 4.2), implemented with lamarckian genetic algorithm, LGA, trough AMDock Software, This oral drug, approved by FDA in 2014, both by Autodock Vina and Autodock 4 has excellent Binding affinity value, ca. −10.00 kcal/mol, a Ki value 40 nM and Ligand efficiency ca −0.15 kcal/mol. These results are comparable to the drug crystallized in the above-mentioned protein, currently used against TBC.

## Author contributions

I.V.F. conceived, designed and wrote the paper, performed the calculations and analysed the data. P.P. overview the article and whore the paper.

